# meaRtools: an R Package for the Analysis of Neuronal Networks Recorded on Microelectrode Arrays

**DOI:** 10.1101/242065

**Authors:** Sahar Gelfman, Quanli Wang, Yi-Fan Lu, Diana Hall, Christopher D. Bostick, Ryan Dhindsa, Matt Halvorsen, K. Melodi McSweeney, Ellesse Cotterill, Tom Edinburgh, Michael A. Beaumont, Wayne N. Frankel, Slavé Petrovski, Andrew S. Allen, Michael J. Boland, David B. Goldstein, Stephen J. Eglen

## Abstract

Here we present an open-source R package ‘meaRtools’ that provides a platform for analyzing neuronal networks recorded on Microelectrode Arrays (MEAs). Cultured neuronal networks monitored with MEAs are now being widely used to characterize *in vitro* models of neurological disorders and to evaluate pharmaceutical compounds. meaRtools provides core algorithms for MEA spike train analysis, feature extraction, statistical analysis and plotting of multiple MEA recordings with multiple genotypes and treatments. meaRtools functionality covers novel solutions for spike train analysis, including algorithms to assess electrode cross-correlation using the spike train tiling coefficient (STTC), mutual information, synchronized bursts and entropy within cultured wells. Also integrated is a solution to account for bursts variability originating from mixed-cell neuronal cultures. The package provides a statistical platform built specifically for MEA data that can combine multiple MEA recordings and compare extracted features between different genetic models or treatments. We demonstrate the utilization of meaRtools to successfully identify epilepsy-like phenotypes in neuronal networks from *Celf4* knockout mice. The package is freely available under the GPL license (GPL>=3) and is updated frequently on the CRAN web-server repository. The package, along with full documentation can be downloaded from: https://cran.r-project.org/web/packages/meaRtools/.

**Author summary:** Cultured neuronal networks are widely used to study and characterize neuronal network activity. Among the many uses of neuronal cultures are the capabilities to evaluate neurotoxicity and the effects of pharmacological compounds on cellular physiology. Multi-well microelectrode arrays (MEAs) can collect high-throughput data from multiple neuronal cultures simultaneously, and thereby make possible hypotheses-driven inquiries into neurobiology and neuropharmacology. The analysis of MEA-derived information presents many computational challenges. High frequency data recorded simultaneously from hundreds of electrodes can be difficult to handle. The need to compare network activity across various drug treatments or genotypes recorded on the same plate from experiments lasting several weeks presents another challenge. These challenges inspired us to develop meaRtools; an MEA data analysis package that contains new methods to characterize network activity patterns, which are illustrated here using examples from a genetic mouse model of epilepsy. Among the highlights of meaRtools are novel algorithms designed to characterize neuronal activity dynamics and network properties such as bursting and synchronization, options to combine multiple recordings and use a robust statistical framework to draw appropriate statistical inferences, and finally data visualizations and plots. In summary, meaRtools provides a platform for the analyses of singular and longitudinal MEA experiments.

## Introduction

The MEA platform is now increasingly being used to study the response of neuronal networks to pharmacological manipulations and the spontaneous activity profiles of neural networks originating from genetic mouse models and derived from human pluripotent stem cells[1-4]. Recent studies aim to not only evaluate wild-type and mutation associated phenotypes, but also to recapitulate the *in vivo* response to various molecules, compounds and drug therapies [5, 6]. Capturing the many and varied activity features from a cultured neuronal network is critical for the full and accurate characterization of that network. However, MEA data are complex to handle. Moreover, an MEA experiment can last several weeks and incorporate many recordings and various treatments. For these reasons, there is a genuine need for methods that can adequately characterize those neuronal networks and also provide valid assessments of phenotypic differences between genotypes and drug treatments in an experiment lasting many days *in vitro* (DIV).

The meaRtools package provides tools to identify complex phenotypes for assessing the effect of mutations and the screening of compounds in a multi-well MEA platform, as presented here in figures 4-6 and as we previously shown [6]. The algorithms described here add to existing methods through calculation of cross-correlation and mutual information between electrodes, as well as enhanced identification of synchronized bursts (including entropy phenotypes for each well). The latter algorithm is shown here to identify recapitulation of *in vivo* epilepsy phenotypes in cultured neurons of the *Celf4* knockout mouse model (*Celf4*^-/-^). Incorporated into the package is also an algorithm that uses electrode-level burst features distributions to identify burst activity variations originating from neuronal subtypes in primary neuronal cultures. An earlier version of the package was recently used to examine the effects of microRNA-128 deficiency on the activity of cortical neural networks [6]. Last, the package provides functions to combine many recordings from multi-DIV experiments and perform rigorous statistical tests of phenotypic differences between genotypes and/or drug treatments.

**Table 1.**
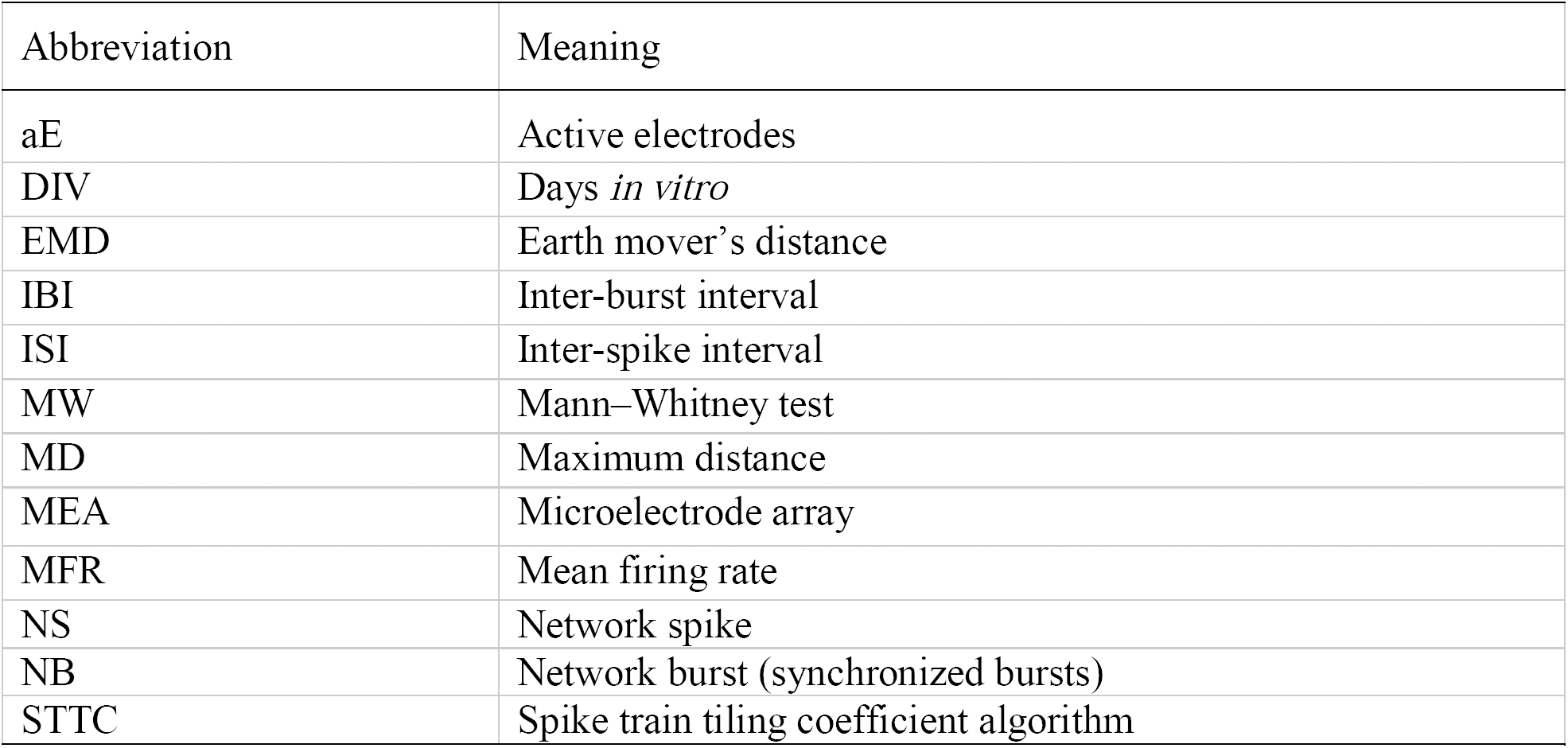
Abbreviations used in text.

## Design and Implementation

meaRtools’s objective is to provide a comprehensive characterization of electrode-level and network activity on a MEA plate, that is composed of one or more wells, each well consisting of multiple electrodes.

The package enables a rigorous examination of differences between various genotypes and/or treatments cultured on the same plate over time. To achieve this purpose, the package provides functions to perform four major analyses (figure 1 and supplementary table S1):

**Figure 1.**
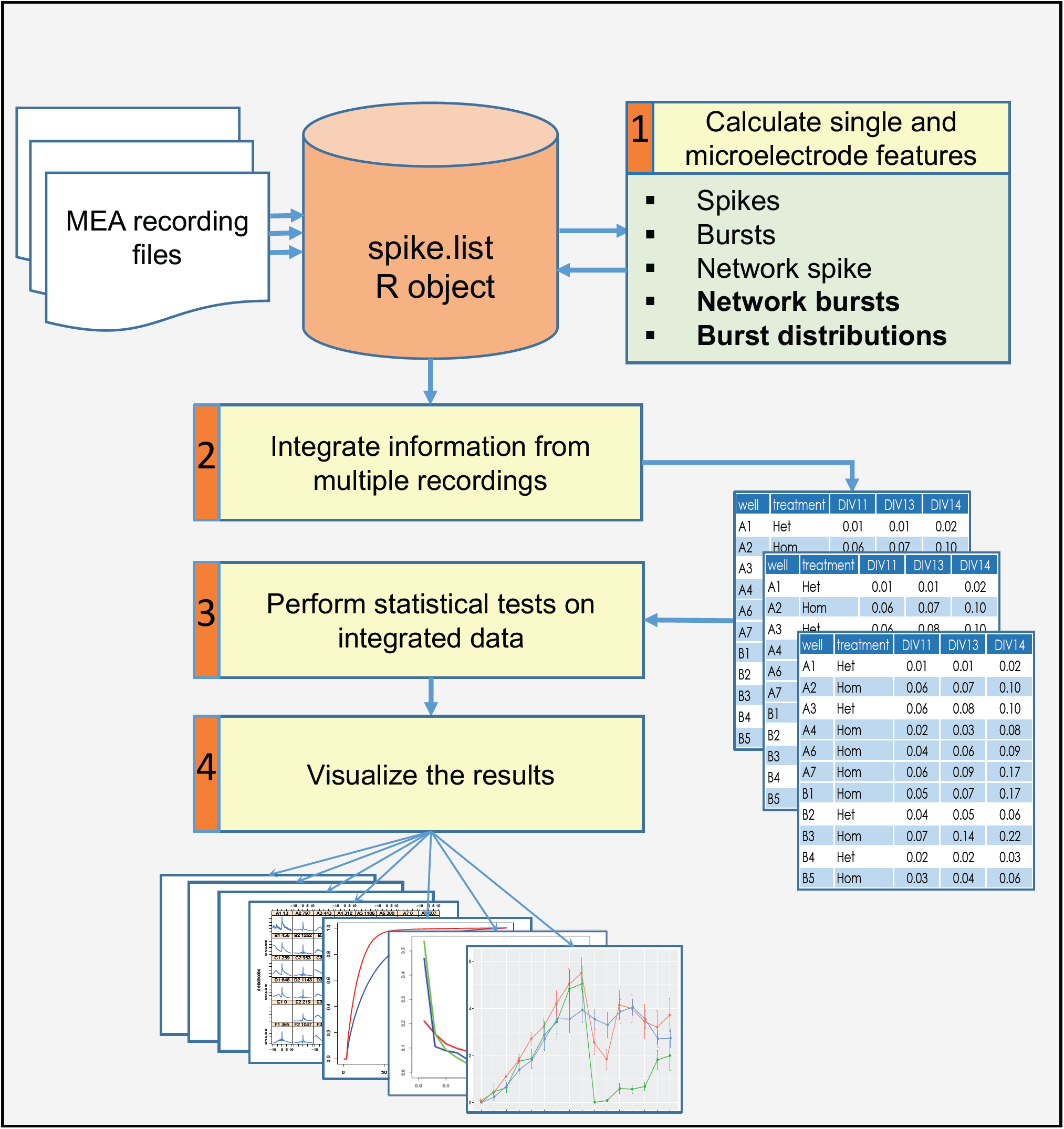
A general scheme of an analysis workflow for several MEA recordings.

1. Identify simple and complex single-electrode and network (multiple electrodes) activity phenotypes. Activity attributes that are extracted include spike and burst features on an electrode-level, as well as features of synchronized network events on the multi-electrode or well-level, such as network spikes (NSs), synchronized network bursts (NBs), cross-correlation and entropy calculations.
2. Combine information from multiple recordings of the same experiment. An MEA experiment can have multiple recordings along several weeks while the neuronal culture is viable. Using meaRtools functions, many recordings of the same plate can be incorporated into a complete dataset, which can be used to test temporal replicability of results with the added advantage of enhanced statistical power.
3. Perform reproducible case/control based statistical analysis. Statistical tests are required when using the MEA platform to characterize and identify differences between genetic models and various treatments. This need intensifies when incorporating data from many recordings and comparing wells grouped by identifiers (e.g. drug or other treatment). The statistical testing scheme presented here is designed to handle this problem specifically.
4. Visualize the results in presentable ready-to-use graphs and charts. The complex picture arising from MEA recordings can be examined on various resolutions: electrode-level activity, well-level synchronization and genotype or treatment activity grouped across several wells and throughout several recordings. These different resolutions can all be visualized using designated functions. Furthermore, comparisons between genotypes and treatments are visualized along with statistical test results.

Since meaRtools is implemented in R, we assume familiarity with the R programming environment and provide a step-by-step workflow for an exemplary experiment consisting of three MEA recordings through the package vignette (https://cran.r-project.org/web/packages/meaRtools/vignettes/meaRtoolsGeneralUsage.html). The vignette provides a thorough guide to meaRtools, beginning with package installation, input handling and the various commands to identify activity features, calculate statistics, and plot. Running the vignette while reading the methods section of this manuscript will provide hands on experience and familiarity with the package. For those new to R, we recommended the introductory material found at: http://www.rstudio.com/online-learning.

### Input and Data Organization

The package can analyze multiple recordings with the same plate layout. The input format is similar to Axion Biosystems ‘spike_list.csv’ format: a comma separated file with a row for each spike holding spike time, electrode name and spike amplitude (mV, supplementary table S2). The input files can be read using the *read_spikelist* function, which then constructs an R object of class ‘spike.list’ that holds the following information retrieved from the input files: electrode names and positions, spike trains (a sequence of action potentials recorded over time) and recording information (start/end time, machine version). The feature extraction functions introduced below add layers to this primary ‘spike.list’ object (supplementary table S1).

The package requires a layout scheme for each experiment that lists all wells used in the experiment and, when applicable, the treatment of each well (supplementary table S3). This layout scheme is necessary to group wells for statistical comparisons of extracted features. A treatment label can represent multiple aspects used to distinguish a specific well, such as a genotype model, a drug treatment exposure, a change in the culture medium, an external stimulus, etc. The term ‘treatment’ will be used henceforth to relate to any of the above.

Although we focus here on analyzing Axion Biosystems datasets, the package can read in data recorded from other platforms, assuming that the data can be manipulated into a straightforward text format. In this case, two files are provided, one containing the spike times recorded on each electrode and one describing the spatial position of each electrode. Our package provides a vignette that demonstrates how such data can be read in and processed; see https://cran.r-project.org/web/packages/meaRtools/vignettes/data_input.html.

### Identifying Simple and Complex Single and Multi-Electrode Activity Phenotypes

The package provides functions for identifying numerous features that characterize various network activity attributes. Features are calculated at three levels: electrode-, well- and treatment-level. Well-level calculations combine the information from all the electrodes within the well. Treatment-level calculations further group wells together by their assigned treatment label.

For the extraction of features for spikes, bursts and NSs described below, the package utilizes code from the open-source Sjemea R package [7] and the algorithms of Eytan and Marom [8] and Legendy and Salcman [9].

### Spikes, bursts and network spikes

Spikes, or single action potentials, are not detected by meaRtools. The package accepts a list of spike time stamps generated using a spike detection algorithm of the user’s choice. To calculate spike activity features and statistics, the package provides the function *calculate_spike_features*, which extracts spike information separately from the spike times of each electrode. Among the extracted features are various spike statistics per electrode and well, such as: 1) The number of active electrodes (aEs) per well, 2) The number of spikes, 3) Mean Firing Rate (MFR, in Hz) and 4) Inter-Spike Interval (ISI), which is the time between two sequential spikes.

Overall, the package provides eleven spiking statistics and features, among which are the well-level Spike Train Tiling Coefficient (STTC), mutual information and entropy measurements discussed below (supplementary table S4).

Bursts are short periods of time with elevated spike frequencies [9]. The package provides an implementation of two algorithms for detecting rapid spiking periods using the function *calculate_burst_features*: the Maximum Interval [10] algorithm and the Poisson Surprise [9] algorithms that have been extensively used previously to identify spike bursts [11-14]. The use of both heuristic and statistical modeling approaches for data comparison allows for an enhanced identification of features [15].

Using either the Maximum Interval or the Poisson Surprise algorithms produces the same burst features, among which are statistics per electrode and per-well for: 1) The number of bursts, 2) Burst durations, 3) Burst rates, 4) Spike rates within bursts, 5) IBIs and 6) ISIs within bursts. Overall, the package provides 19 burst features (supplementary table S4).

Network spikes (NSs) are synchronized events of neuronal populations in a short period of time. The meaRtools package detects these events using a well-established algorithm developed by Eytan and Marom [8] that was obtained from the sjemea package [7] and previously used for characterizing synchronized activity [16, 17]. The function *calculate_network_spikes* detects NS within each well. A NS is detected when there are at least a user-defined number of aEs (default 4 aEs, or 25% of total number of electrodes) detected within a user-defined time window (default 10 ms).

NS features are next extracted using the function *summarize.network.spikes,* and include basic statistics for: 1) NS number, 2) ISIs in NS, 3) number and 4) percentage of spikes participating in NS. Overall, the package provides a total of ten NS features (supplementary table S4).

### Algorithms to assess network synchronization and burst activity patterns

#### Spike Train Tiling Coefficient

We have implemented STTC to evaluate pairwise correlations between a pair of electrodes [18]. STTC has recently shown to examine the effects of an agonist of acetylcholine receptor and several inhibitory antagonists on network synchronization [19, 20].

We implemented STTC as follows: Given *N* active electrodes within a well, we ignore any possible dependence of distance upon correlation and simply calculate the average STTC from all *N*(*N*-1)/2 pairwise correlations.

Average well-level STTC values per well can be computed using the *compute_mean_sttc_by_well* function.

#### Entropy and mutual information

Entropy and mutual information have been used previously to describe and characterize neural spike trains [21-23]. We adapted the two information theory metrics based on the original ideas presented by Shannon [24] to MEA data analysis. These metrics have utility for characterizing recordings of a single well, a set of wells per treatment, and comparing treatments from the same plate.

Entropy was used as a broad measurement of the amount of disorder measured at an MEA electrode, as well as across a well. For entropy calculations, we used the standard equation for calculating the entropy of a system:

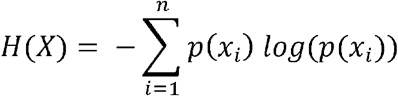

We define H(X) as the calculated entropy measurement for electrode X. i represents a time interval bin, out of *n* total separate equally sized time interval bins (default is 0.1s bin size) from start to end of the recording time. In the probability distribution, Xi is set as the number of spikes in the i’th bin of electrode X divided by the total number of spikes observed in the full recording. For a well we use a mean statistic across all electrodes within well. When testing a full plate, we use the mean entropy across a group of wells (combined by treatment) as a test statistic representing the ‘orderliness’ of the firing patterns. Sets of mean entropies per treatment can be later compared to determine if there is evidence of a shift in the distribution between two different treatments. Normalized entropy statistics can be collected per well with the *calculate_entropy_and_mi* function.

The second metric, mutual information, can be used to compare patterns from two separate electrodes, and could be extended to represent the network level activity of neuronal firing in a particular well. This method was used to identify differences in synchronized activity between neuronal networks from the *Celf4*^-/-^mouse model of epilepsy and wild-type (*Celf4*^+/+^) control neuronal networks (see Results). We start with the generalized equation for mutual information (MI):

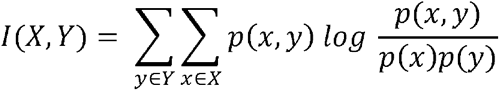

We define I(X,Y) as the information shared between electrodes X and Y. We define a number of equally distributed time interval bins in the time period and for each electrode, count the spikes in each b_i_. As before, we transform X and Y into separate probability mass functions, where the probability of a spike falling in a particular time interval bin equals the count at that bin divided by the total number of spikes that detected in the recording.

Given that the number of spikes observed during recording time can vary between electrodes, it was important to further transform X and Y to take this into account. For this we take the spike count in a time interval to infer the presence or absence of a burst, and as such we can classify each time interval at an electrode as either a burst member or non-member. To do this we transformed each input vector X such that the value X_i_ equals 1 if the spike count is greater than the 75th percentile of spike counts across all bins in X, and set as 0 otherwise. Such a simple transformation of the transformation of the data means that the probability mass function for X is collapsed down to p(X=0) and p(X=1). Subsequent MI calculations are far more efficient since in terms of combinatorics, only 4 outcomes at a given time interval are possible: X=0/Y=0, X=1/Y=0, X=0/Y=1 and X=1/Y=1. As such we explicitly compute the mutual information between electrodes X and Y as:

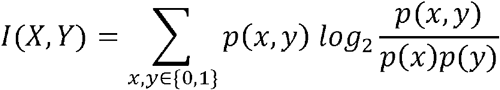

We produce a distribution of these pairwise statistics per well, and aggregate the statistics per treatment. A Mann–Whitney (MW) test can be later performed on these sets of values between two treatments to determine if there is evidence for neurons in one treatment having a higher level of coordinated network level firing than in another treatment. Average well-level pairwise mutual information values per well can be computed using the *calculate_entropy_and_mi* function.

#### Burst features distributions

Distributions of burst features can be compared and visualized to assess certain burst features by looking at their density distributions along a recording. This method was previously used to successfully test the effects of a miR-128 knockdown in cultured neuronal networks [6], and was used in this study to discern between various culture treatments and cell density (see Results). The reason behind constructing this method is that primary cultures contain multiple neuronal subtypes (e.g. GABAergic and glutamatergic), which demonstrate different activity signatures [25, 26]. Thus, merely extracting a mean and standard deviation of a feature will misrepresent the activity fluctuations that arise from the combined activity of neuronal subtypes in the network. For example, spike frequencies within bursts may differ between GABAergic interneurons and glutamatergic neurons [27], with GABAergic neurons often exhibiting narrower spike wave-forms and faster-spike activity [28, 29]. Furthermore, certain anti-epileptic or anti-psychotic drugs selectively target specific neuronal subtypes. However, this selective effect may not be observed when comparing the average change of an entire cultured network.

The function *calc_burst_distributions* calculates empirical distributions for bursting features and compares them between treatments using two independent methods (figure 2A, see Statistical Testing and Visualization).

**Figure 2.**
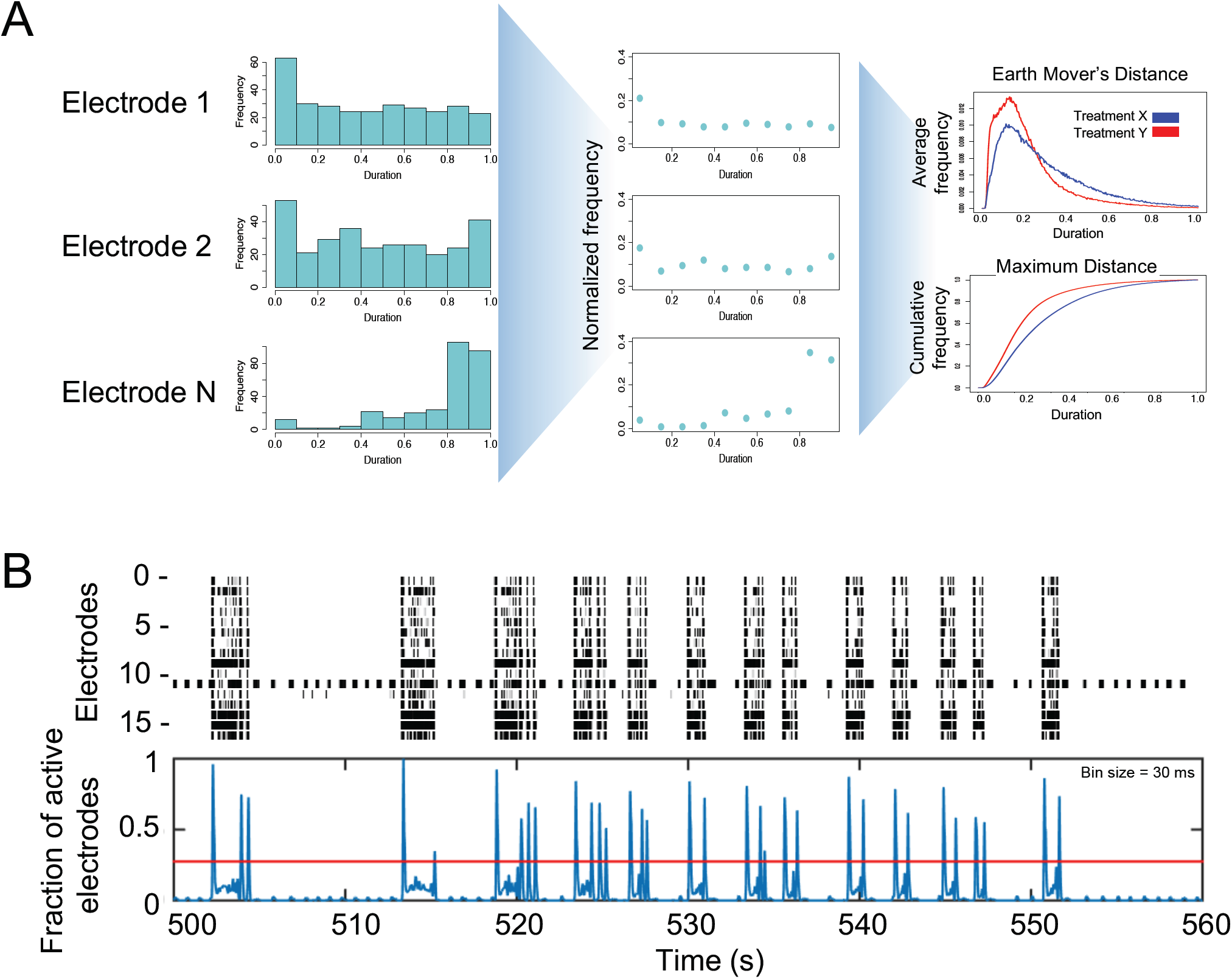
Schemes for computing NBs and burst distributions. **A)** Creating burst features distributions. First, burst feature histogram is calculated for each electrode (left panel). In this example, it is calculated for burst duration. Next, histograms are normalized to number of values, resulting in a 0-1 value. Last, all electrodes are averaged to create a normalized distribution plot (top right panel) and a cumulative plot (bottom right panel) for each tested treatment. **B)** Detecting synchronized bursts. Spike data from a raster plot (upper panel) showing the spikes (x-axis) for each active electrode (y-axis) is binned and combined through a weighted Gaussian kernel smoothing method to generate the fraction of active electrodes (blue lines in lower panel). The Otsu global thresholding algorithm [32] is then applied to identify intervals above the threshold (red horizontal line) as synchronized NBs.

Density distributions are calculated for five burst features: IBIs, ISIs within bursts, number of spikes in bursts, burst durations and spike frequencies within bursts (firing rate, Hz). For each feature, the algorithm adjusts for variability between electrodes in a well. This is done by calculating the histogram of a feature in each electrode separately (figure 2A, left panel) and normalizing it to values between 0-1 (figure 2A, middle panel). Next, all normalized histograms are grouped and averaged by treatment labels. The algorithm permits performing this step also by grouping electrodes first by wells and then averaging well information by treatment. To later test for differences between treatments, the package provides a function which performs distribution comparison tests, permutes electrode labels and plots the results (figure 2A, right panel, see Statistical Testing and Visualization).

#### Synchronized network bursts

We consider NBs as bursts appearing at several electrodes simultaneously, that are longer and more intense synchronization events than NS and correspond to electrode-level burst activity that is synchronized across electrodes in a well. The underlying reason to identify NBs is that, while the NS detection algorithm identifies short network synchronized activity lasting tens of milliseconds, synchronized bursting events were shown to last tenths of seconds to seconds in MEA experiments [15, 30, 31]. To catch these long synchronized network events, a method was constructed that investigates bursting patterns within wells and also between wells clustered based on treatments.

The function *calculate_network_bursts* combines burst information at the electrode-level into well-level data as the presentation of synchronized bursts across a well (figure 2B). First, spike time within spike trains from all electrodes is binned using a bin size of 2ms to guarantee that at most one spike is called within each bin. Next, a Gaussian filter with user-defined window sizes (defaults are: 10, 20 and 50 ms) is applied to smooth the binned spike trains from each electrode. The smoothed signal is then further standardized to have a maximum signal value of 1. All smoothed signals at the electrode-level are then combined and smoothed again using the same Gaussian filter. The final result from this step is a smoothed signal at each given window size that measures the overall synchronization of all electrodes in a well, with larger values indicating higher level of synchronized bursting activities. Then, the Otsu global thresholding method is applied to the well-level signal to automatically detect burst intervals [32]. This method was chosen for its simplicity and parameter free nature, although other methods, such as adaptive thresholding, can be utilized. Last, based on the NB intervals obtained from Otsu thresholding, NB information is collected at the well-level.

The algorithm extracts statistics for: the number and rate of NBs, the number and percentage of spikes participating in NBs and the spike intensities within NBs, which is the spike rate within NBs. These NB statistics were previously shown to infer the biological effects of miR-128 knockdown in cultured neuronal networks [6] and are shown in this work to successfully identify significantly higher synchronization of bursts in a mouse epilepsy model (figure 4B-4C) and in the homozygous *Celf4*^*-/-*^ mouse model (figure 6B-6C).

**Figure 3.**
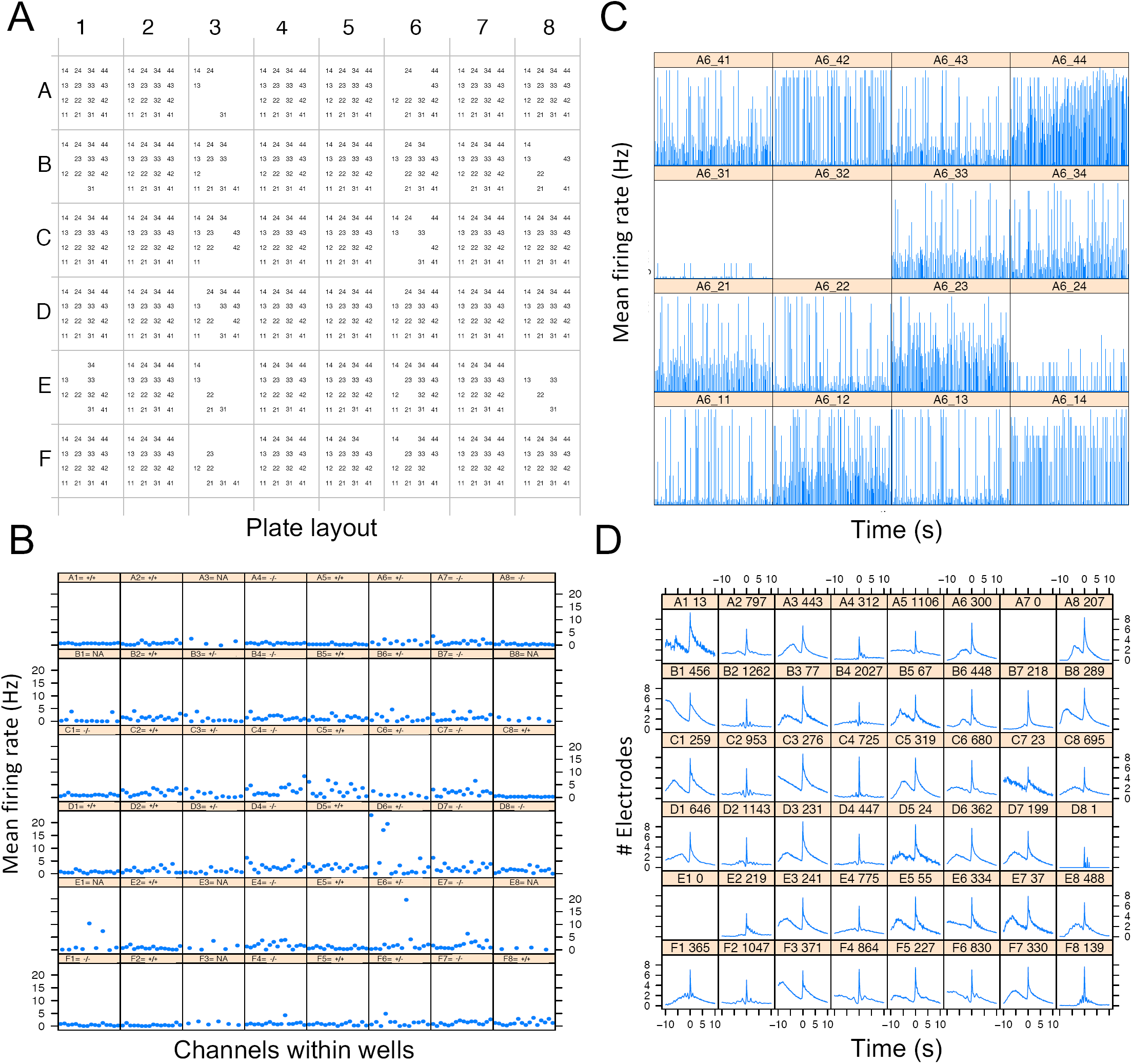
General information of plate activity. **A)** A matrix representing a 48-well plate, where each well consists of 16 electrodes. aEs are represented by name consisting of column+row position in each well (“11” for first electrode to “44” for the last). **B)** MFR (Hz) for aEs per well in a 48-well plate. Title for each well has the genotype label of the well: +/+ (wild-type), −/− (homozygous), +/− (heterozygous), NA (Not available). **C)** Average MFR of all 16 electrodes of well A6, presented for each second of a 900s recording. **D)** Average number of electrodes participating in NSs around the peak of a network event. The x-axis represents user-defined time bins (default is 100 ms) before and after a NS peak (−10 equals 1s before the peak). Title for each well consists of well-name and number of identified NSs.

**Figure 4.**
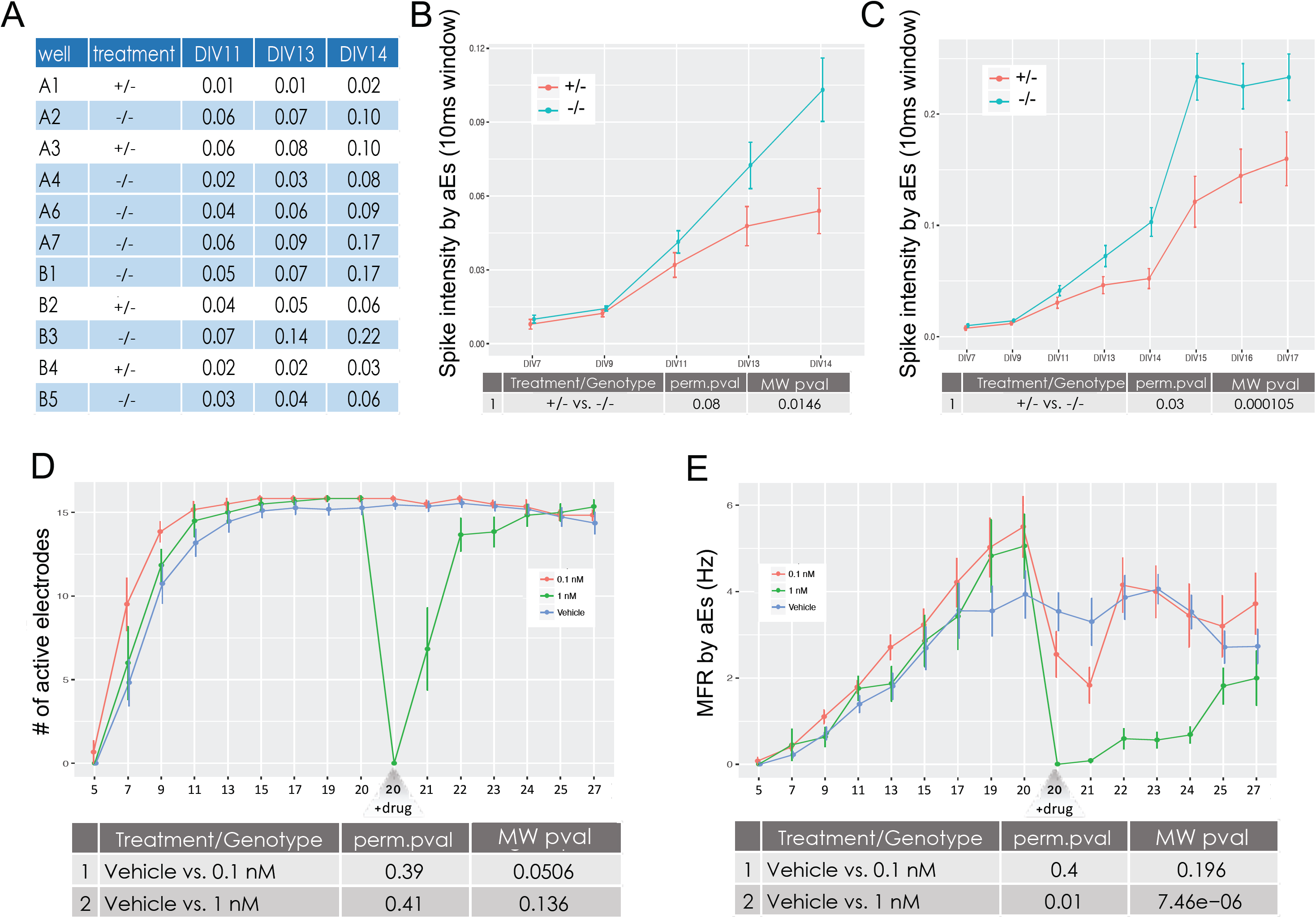
Comparing features between treatments. In panels A-C, treatment refers to genotype (heterozygous, +/- and homozygous, -/-). In panels D and E, treatment refers to drug treatment. **A)** Output tables are printed for every feature, holding average values (in this case, Spike Intensity per aEs) per well (rows) and for every recording analyzed (columns). **B)** Spike Intensity per aEs for two genotypes by DIV on a subset of five recordings. This graph corresponds to the full aggregated table of A). **C)** Same as B), but for a subset of eight recordings. **D)** Comparing the number of aEs for wells treated with a vehicle or one of two channel blocker treatments (0.1nM and 1nM) along a 27 DIV experiment. **E)** MFR for the same experiment as in D), showing differences in the effects of drug concentrations.

**Figure 5.**
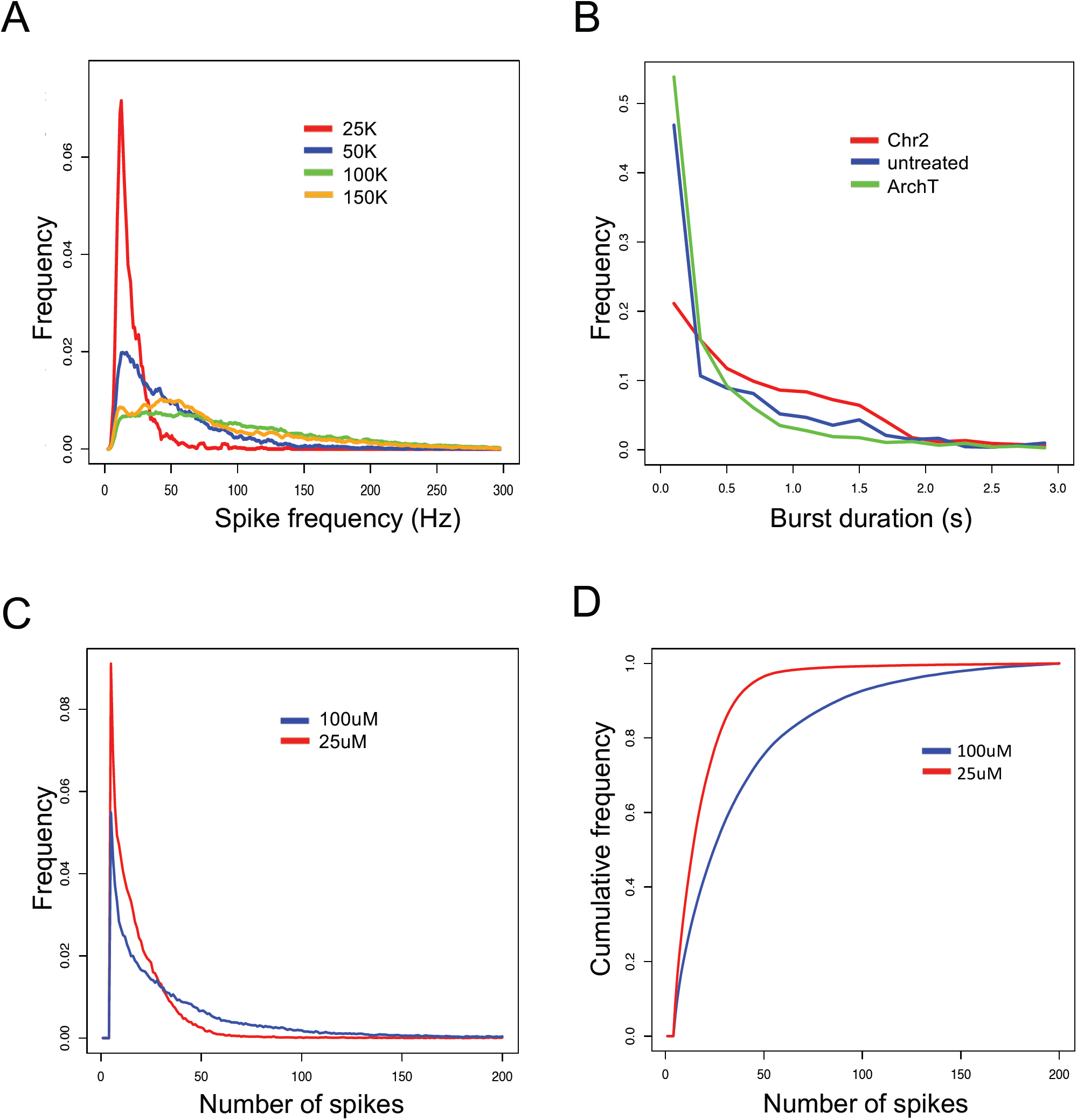
Burst features distributions. **A)** Frequencies (y-axis) of spike-rate in bursts (x-axis) are calculated for four different cell density cultures in a single recording. A user defined maximum of 300 Hz is set. **B)** Frequencies (y-axis) of burst durations (x-axis) are presented when introducing two different microbial light sensitive membrane proteins vs. untreated cells in a single recording. A user defined maximum for burst duration was set to 3 seconds. **C)** Combining burst features distributions of number of spikes in bursts from 24 sequential same-plate recordings. Treatments were automatically tested for difference using the EMD test and results were displayed after permutations. **D)** Cumulative distributions of the same data as in C), treatments were automatically tested for difference using the MD test and results were displayed after permutations.

**Figure 6.**
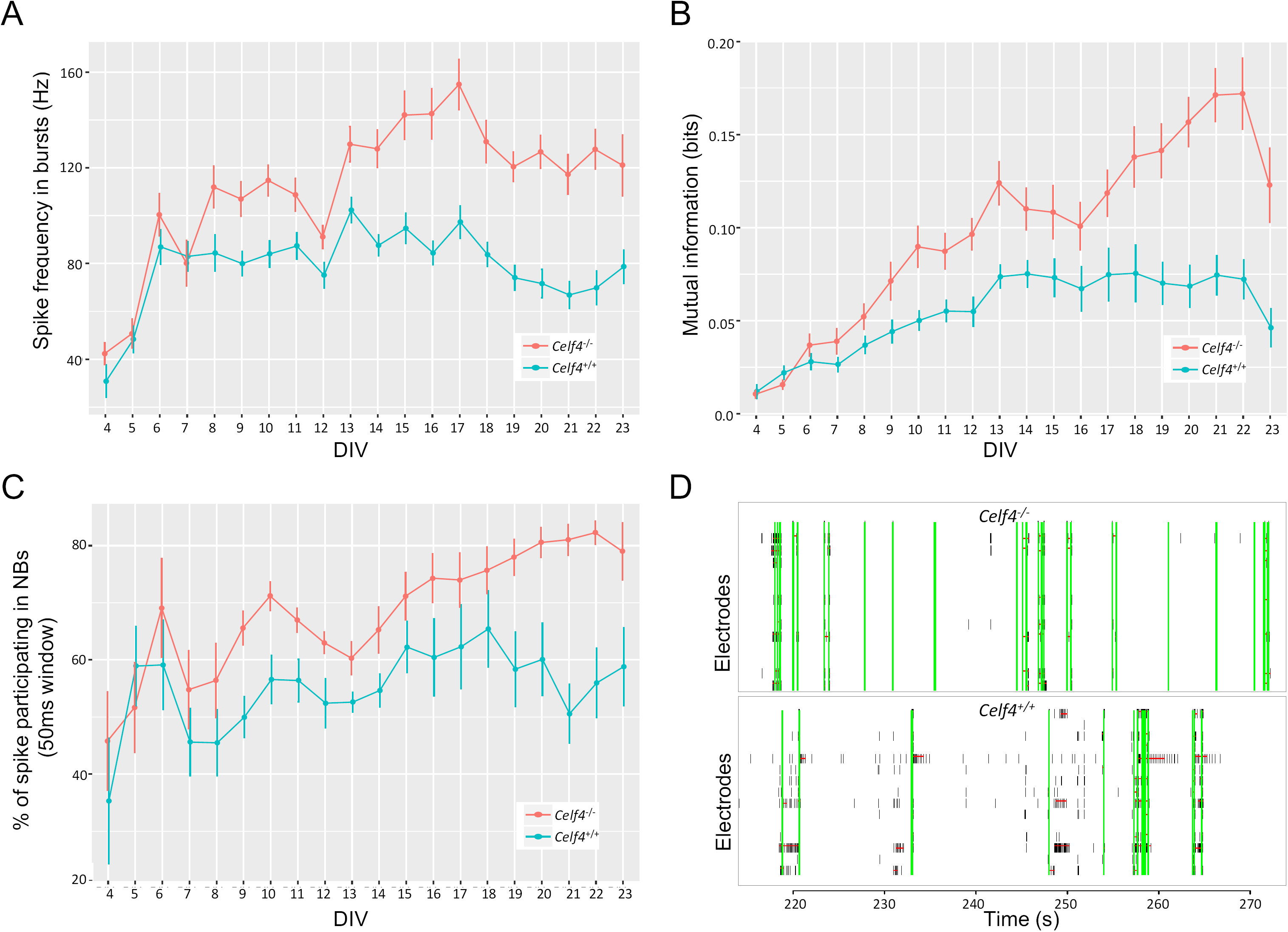
*Celf4*^-/-^ neurons show elevated network synchronization phenotypes. **A)** Spike frequency in bursts for *Celf4*^+/+^ (blue) and *Celf4*^-/-^ (red) neural networks. **B)** Mutual information between electrodes is increased in *Celf4*^-/-^ networks (red). **C)** The percent of spikes participating in NBs is increased in *Celf4*^-/-^ neurons (red), suggesting that ratio of spikes that participate in synchronized network events is higher in the *Celf4*^-/-^ networks. **D)** Raster plots show NSs (vertical green lines) and bursts (horizontal red lines) for two adjacent wells: a *Celf4*^-/-^ well (upper panel) and a *Celf4*^+/+^ well (lower panel). *Celf4*^-/-^ exhibits few sporadic spikes between highly synchronized events, while *Celf4*^+/+^ exhibits less network events and more sporadic spikes. Raster plots present 60 seconds of 900 seconds recordings.

Overall, the package provides 11 NB features for each time window to a total of 33 features (supplementary table S4).

### Combining Multiple Recordings

MEA experiments are constructed from multiple recordings of the same plate over a certain period of time. Correctly assessing the activity and differences over time requires analyzing several recordings as one set of information with various time points. The package provides functions for combining several recordings and filtering wells from the combined dataset based on inactivity measurements. All feature extraction functions store the extracted information in the same ‘spike.list’ object. The function *aggregate_features* uses the information stored in the ‘spike.list’ object to combine data from all the analyzed recordings into an aggregated table for each feature. The aggregated tables have recording labels as columns and well-labels as rows, and can be printed as csv files or used later for treatment comparisons. The package also provides the function *filter_wells* to exclude inactive wells from these aggregated tables. An active well is measured using a minimum number of aEs (default 4, or ¼ of the total number of electrodes). The function *filter_wells* considers whether a well has been active in more than a certain percentage of recordings in the experiment (default is 50%). Inactive wells that fail to meet this criterion are not used when comparing treatments.

### Statistical Testing and Visualization

The combined tables from multiple recordings can next be used to compare treatments along the experiment. The function *permute_features_and_plot* performs all the necessary statistical tests between treatment labels and plots the results of all features in .pdf format in a designated output directory. The tests are performed as follows: First, a MW test is performed to compare distributions of each feature between treatments, and the resultant p-value is recorded. Next, a permutation scheme is performed where the treatment labels of the active wells are randomly shuffled X times (default 100) while the observations within each well are kept intact. This preserves correlations between time points within wells while breaking any relationship with treatment and subsequent outcome. A permutation p-value for the original MW test is computed as the proportion of permuted data MW p-values that were less than or equal to the MW p-value of the original un-permuted dataset. Last, for each feature, a graph and a table (csv format) are printed with the mean and standard error (SEM) of the measured features for each of the recordings.

Comparing and plotting burst distributions, is done using the function *dist_perm*. The algorithm works as follows: For each of the five burst features, the function *calc_burst_distributions* generates a normalized histogram per electrode (supplementary table S5). Next, the function *dist_perm* groups all distributions by treatment labels and compares them between treatments using two methods: 1) The Earth Mover’s Distance (EMD), using R package emdist [33] and 2) the Maximum Distance (MD) between the cumulative distributions of the normalized histograms.

Once the test results for EMD and MD are computed for the original dataset, a permutation scheme is performed where the treatment labels of the active wells are randomly shuffled X times (default 100) while the distributions within each well are kept intact. A permutation p-value for EMD is then computed as the proportion of permuted data EMD values that are equal to or greater than the original EMD value from the un-permuted dataset. A permutation p-value for MD is defined similarly. Last, both the normalized and cumulative histograms are plotted with the final permuted p-value.

#### Performance

The performance of the package was tested by analyzing three recordings included in the package as example files recorded for 40s, 60s and 60s on a 48-well plate with 16 electrodes in each well. We ran the full package vignette, extracting all possible features and statistics for each recording including: spike statistics, mutual information, STTC, burst statistics and distributions, NSs and synchronized NBs. The test also included combining data statistics from all recordings of the experiment and performing 100 default permutations for testing differences between three available treatments for all 73 available features. The test concludes by plotting the comparison between treatments for all the features tested. On a MacBook Pro with a 2.8 GHz Intel Core i7 CPU and 16 GB of 1600 MHz RAM, the average execution time for this test was 3.23 min and 4.97 min, excluding or including permutation and plotting, respectively. Increasing the number of permutations to 1000 increased the run time significantly to 16.05 minutes.

## Results

The meaRtools package can be used as an analysis pipeline on an experiment composed of data from several recordings and with several treatments. The package is shown here to extract novel biological insights by identifying neuronal phenotypes of a *Celf4* epilepsy mouse model [34] and was also recently utilized to assessing the effects of miRNA deficiency on neuronal activity [6], both of which demonstrate the ability of the algorithms presented here to identify and provide robust statistical measurements for simple and complex network-associated phenotypes. Below, we present various experimental examples to illustrate the diverse features and uses of meaRtools.

### General Activity Information

To gain a preliminary view of plate activity, the package generates graphs of activity measurements for three levels of data: electrode-, well- and plate-levels. Presented here are a subset of these graphs for a single recording in a 48-well plate (16 electrodes per well) containing cultured cortical neural networks from the brains of postnatal wild-type mice. The lowest resolution map of the plate shows a matrix of all aEs for each well in the plate (figure 3A). A higher resolution graph shows electrode activity per well as the MFR of all aEs (figure 3B). Even higher resolution shows the MFR of each electrode in each well (figure 3C represents a 900s recording). The latter is plotted for each well separately.

In order to demonstrate network behavior before and after each NS, we show the number of electrodes participating in NS around the peak of a network event (figure 3D). For example: some networks present a decline in participating electrodes before the event takes place (wells B1, C1 and D1 in figure 3D), while others exhibit a gradually increasing number of electrodes participating in a NS before a fast accumulation of electrodes leading to the NS peak (wells F1 and F8, figure 3D).

The full set of graphs that can be printed by the package is available as supplementary information and includes log ISI statistics, spike statistics within bursts and other network information. These graphs provide an overall view of the activity of a single recording.

### Longitudinal Modulation of Network Activity by Drug Treatment

For every extracted feature a comparison can be made between treatments in a multiple-recording experiment. For every activity attribute (i.e. spikes, bursts, NSs and NBs), an output folder is created with .csv files for every feature, holding average values per well for every recording analyzed. An example is shown here for Spike Intensity within NBs representing DIV 11-14 analysis of two genotypes of a genetic mouse model of epilepsy; which are heterozygous (+/-) or homozygous (-/-) for a specific mutation (figure 4A). Also printed are graphs showing mean and standard error for each feature. These graphs compare the treatments over all the analyzed recordings (figure 4B-4E). The output directories include tables and graphs for a total of 70 features: eight spike features, 19 burst features, 10 NS features and 33 NB (see supplementary table S4).

The comparison analysis is flexible and can be performed for subsets of recordings and treatments. The example in figure 4B and 4C shows a comparison of network activity between the two genotypes (+/- and -/-). Under each graph are the results of multiple MW and permutation tests performed between the genotypes. The Spike Intensity within Network Bursts feature (spikes per NB per sec) analyzed here illustrates the synchronization level of a network. A comparison of treatments, using five recordings from an experiment comprised of 14 DIV, shows a trend for higher spike intensities in the homozygous genotype, which is not significant after a permutation test (figure 4B). However, inclusion of data from three more DIV shows a significantly increased network synchronization in the homozygous genotype (figure 4C).

The feature comparison analysis is not limited to the number of recordings or treatments. In figure 4D, the number of aEs are shown for an experiment spanning 27 DIV. This experiment had three treatments: a vehicle treated control and two concentrations (0.1nM and 1nM) of a sodium channel blocker (figure 4D-4E). The treatment was administered on DIV 20 and the plate was recorded for 7 DIV following drug administration. Analysis of the number of aEs shows no significant difference between groups before the drug was added, and a dramatic decrease in number of aEs in the wells treated with 1nM but not those treated with 0.1nM (figure 4D, green vs red lines, respectively). As expected, MFR analysis indicates a significant decrease in MFR for both 0.1nM and 1nM drug treatments[35]. Moreover, the kinetics of MFR decrease are relative to the drug concentration, as is the time it takes the MFR to return to untreated values (figure 4E, red and green lines).

Overall, these results present the flexibility of the functions to handle varying number of recordings and provide true biological insights. The ability to test treatment differences over many recordings, presented here for the first time, provides a strong and valuable tool to assure drug, compound or genotype effect.

### Discerning Various Culture Treatments Using Burst Feature Distributions

In addition to the comprehensive examination of synchronization between electrodes, meaRtools provides a way to identify changes in bursting activity characteristics throughout the recording through examining distributions of burst features such as duration and spike frequency within bursts. Burst features distributions can account for the differing behaviors of neuronal subtypes in a primary culture. For example, burst activity might have differing durations, fluctuating spike rates and other features that are influenced by different activity profiles of specific cell types. While these varying activity properties might not be caught using simple statistical measurements, they can be identified using empirical distributions of features.

Burst features distributions can be compared between treatments in single or multiple recordings. Single recording treatment comparisons are tested using the Kolmogorov-Smirnov test (K-S test) for comparing probability distributions. For instance, when comparing the effects of cell density on network behavior, a significantly higher proportion of low Spike Frequencies is observed at low cell density of 25,000 (25k) cells relative to higher densities (figure 5A, red line), suggesting that higher cell densities have higher spike frequencies within bursts. In a separate experiment, we compared the effects of two different microbial light sensitive membrane proteins: Channelrhodopsin-2 (Chr2) and Archaerhodopsin-T (ArchT) on burst duration. We observed a higher ratio of long burst durations for ‘Chr2’ that is not significant between treatments (figure 5B, red line).

Comparisons of the distributions of burst features can be performed over multiple recordings. The analysis of number of spikes in a burst is presented here for an experiment with 24 recordings that were combined using the methods above. The differences between treatments are calculated using two measurements of distances (figure 5C-D), and followed by permutation tests. This specific analysis successfully validates that a treatment using a specific compound treatment at 25μM (figure 5C-D, red line) has significantly more bursts with low number of spikes than the 100μM treatment (blue line).

### Identification of excitability phenotypes in a mouse seizure model

We utilized the full capabilities of meaRtools to identify and compare complex activity phenotypes of a *Celf4*^*-/-*^ mouse seizure model. [34, 36]. Here, we demonstrate that the MEA platform, analyzed with meaRtools, can identify epilepsy-like phenotypes in neuronal networks from *Celf4*^*-/-*^ mice.

Seizures are often characterized by hypersynchronous discharges that may occur at a specific region of the cortex and spread into neighbouring brain areas. The cellular mechanism of seizure initiation is thought to be the network hyper-synchronization and high frequency bursts consisting of increased density of action potentials, presumably due to an excitation/inhibition imbalance [37]. Sufficiently synchronized bursts may pass the threshold of surrounding inhibition and activate neighboring neurons leading to broader recruitment, network propagation and ultimately seizures. While the *in vitro* manifestations of seizures are not fully understood, it is thought that both increased synchronicity of network firing and increased bursting are analogous to the *in vivo* phenotype [38]. As mentioned, *Celf4* deficiency is known to cause neurological phenotypes in mice including, most prominently convulsive seizures [34, 36]. Here, we use *Celf4*^*-/-*^ mice as “proof of concept” for whether meaRtools synchronization algorithms can identify aberrant excitability phenotypes that are often characteristic of mouse seizure models.

We examined the various features and statistics that meaRtools computes and found that *Celf4*^*-/-*^ neurons consistently showed significant elevation in several features compared to wild-type (*Celf4*^*+/+*^) neurons, among which were: increased frequency of spikes in bursts (permutation p value < 0.01), increased mutual information between electrodes (permutation p value < 0.01) and an increase in various NB phenotypes (figure 6, permutation p values < 0.01 - 0.02, supplementary table S6). Together, these features offer a view of the change in network synchronization of *Celf4*^*-/-*^ mice, and point to a higher density of spikes in bursts and a higher synchronization of bursts between neurons belonging to the same neuronal network. This phenomenon was tested across an experiment lasting 23 DIV, where *Celf4*^*+/+*^ and *Celf4*^*-/-*^ genotypes were cultured on 12 and 11 active wells, respectively. The final statistical tests were performed on recordings from DIV 15-23, starting when the networks were developed and reached a stable MFR and ending with cell death. The results were replicated in another experiment lasting 23 DIV, where each genotype was cultured in nine wells.

We have also examined the change in MFR between the genotypes, and find a slight increase in MFR in the *Celf4*^*-/-*^ neurons (supplementary Table S6). However, this increase did not survive the meaRtools permutation scheme and could not explain the significant increase in synchronization phenotypes observed in figure 6.

## Discussion

The package presented here was constructed with the intention of providing a platform with a wide range of MEA analysis capabilities. We made sure it incorporates commonly used algorithms for spike activity analysis with new capabilities, presented in meaRtools for the first time, such as the ability to combine MEA recordings and apply rigorous statistical tests to compare between groups of wells with varying treatments.

Included in meaRtools are methods for burst and activity measurements that are currently available in commercial software packages, such as NeuroExplorer [10]. Features that are common to both tools are routinely used for network analysis and include standard spike and burst metrics, such as firing rate, number of bursts detected, percentage of total spikes that are contained in bursts, inter-spike and inter-burst intervals, etc. Furthermore, the maximum interval burst detection algorithm is implemented based on the algorithm presented in NeuroExplorer. Activity features we report here that are unique to meaRtools include additional synchrony measures, such as the spike train tiling coefficient (STTC), entropy and mutual information and synchronized NBs analysis. Furthermore, meaRtools is a freely available open-source package that leverages the powerful statistical analysis capabilities and flexibility of the R programming language to provide a consolidated comparison of multiple experiments into a single, summarized statistical and graphical output. Lastly, although this paper describes meaRtools in conjunction with *in vitro* MEA analysis, the algorithms presented here can be equally applicable to *in vivo* MEA work, and in general to the analysis of any action potential spike time stamp data.

In conclusion, the meaRtools package extracts a detailed report of over 70 activity phenotypes (features), both previously existing and novel. The package provides a single platform to handle multiple recordings of the same experiment, and the tools to perform statistical comparisons between treatments on these multiple recording experiments.

## Availability and Future Directions

The meaRtools package is open-source and freely available under the General Public License version 3.0 (GPL>=3). The package is available on the CRAN web-server repository (https://cran.r-project.org/web/packages/meaRtools/index.html). Updated source-code can be found at https://github.com/igm-team/meaRtools. The package provides a step by step vignette for running an MEA analysis pipeline using exemplary datasets in an effort to make MEA analysis accessible to all.

Here we explain the major analyses that can be done using meaRtools and focus on several features to perform detection of phenotypic differences. However, the current version of the package detects 73 features and five feature distributions that can be used in the holistic evaluation of an MEA experiment. Users are encouraged to explore all the features and capabilities the package entails with the help of the package vignette.

The package is updated regularly; each version incorporates additional capabilities to detect and test additional phenotypes. Current work focuses on adding machine learning algorithms to distinguish between treatments, graphical representation of NBs and activity pattern recognition algorithms within and between wells.

